# Homophilic interaction of E-Cadherin prevents cell-cell fusion between developing germline and surrounding epithelia in *Drosophila* ovary

**DOI:** 10.1101/2022.11.22.517537

**Authors:** Matthew Antel, Rachel Norris, Mayu Inaba

**Affiliations:** Department of Cell Biology, University of Connecticut Health, Farmington, CT 06030

## Abstract

In the *Drosophila* ovary, developing germline cysts are encapsulated by somatic follicle cell epithelia and E-Cadherin localizes to the interface of these tissues. E-Cadherin mutants have been shown to have multiple defects in oogenesis. Therefore, it is difficult to determine E-Cadherin function on germline-soma interaction. In this study, we characterize E-Cadherin function, specifically focusing on germline-soma interaction. Unexpectedly, knockdown of E-Cadherin either in the germline or follicle cells results in excess formation of membrane protrusions at the interface of these cells, which leads to a cell-cell fusion and indicates that homophilic interaction of E-Cadherin is required for maintenance of the tissue boundary between these two adjacent tissues. The fate of follicle cells fused to the germline becomes compromised, leading to a defective individualization of germline cysts. We propose that homophilic interaction of E-Cadherin facilitates a barrier between adjacent tissues, demonstrating a unique model of cell-fate disturbance caused by cell-cell fusion.

## Introduction

The *Drosophila* ovary serves as a particularly useful model for the study of how multiple cell types interact and coordinate with each other to complete successful organ development. Each ovary consists of 16-20 ovarioles, structures that contain the oocyte and somatic support cells at various stages of development [1–3]. The tip of the ovariole is the germarium, which houses the stem cells of both germ and somatic lineages [1–5]. In the niche, 2-3 germline stem cells (GSCs) adhere to somatic niche cells, called cap cells, by adherens junctions [6]. The GSCs divide asymmetrically to produce cystoblasts (CBs) which contact escort cells (ECs) and divide mitotically to produce a cyst of 16 interconnected germ cells which transit through the anterior compartment of the germarium and contact follicle cells (FCs) [1–3]. The FCs encapsulate the germline cyst, forming a simple epithelium around the germ cells [4, 5]. One of the 16 germ cells within the cyst is specified as the oocyte, and the remaining 15 germ cells become nurse cells (NCs). The FC-encapsulated cyst, now called an egg chamber, exits the germarium and undergoes egg chamber development [1–3, 7]. Throughout egg chamber development, the epithelial FCs and underlying germ cells maintain a close association to one another which facilitates growth and successful development of the oocyte [1–3, 7].

E-Cadherin (called shotgun/shg in *Drosophila*) serves as a calcium-dependent, homophilic cell–cell adhesion receptor [8, 9]. E-Cadherin is involved in multiple biological processes, including establishing and maintaining cell adhesion [10, 11], cell migration [12] and spindle orientation [13]. E-Cadherin is a major component of adherens junctions which have been characterized in many *Drosophila* tissues and are essential for the entire life cycle, including within the embryo where they are indispensable for morphogenesis [10, 14, 15]. Within the adult male and female germline, adherens junctions adhere germline stem cells to their niche and thus are essential for germline development [6, 13]. Adherens junctions can also be found adhering neighboring FCs. These FC-FC junctions are cadherin-dependent and required for the proper morphogenesis of the FC epithelium [16, 17].

E-Cadherin is essential for later stages of oogenesis. Developing germline cysts are composed of one oocyte and 15 nurse cells (NCs). Oocyte growth is supported by cytoplasmic connections with NCs through intercellular bridges, called ring canals [18], and E-Cadherin is required to anchor ring canals to FC’s plasma membrane to support growth [19]. Moreover, E-Cadherin-based adhesion between the oocyte and FCs regulate oocyte positioning and egg chamber polarity [17, 20, 21].

E-Cadherin has been shown to localize to the interface of NCs and FC epithelia [12]. However, it is unclear whether E-Cadherin is involved in interactions between these cell types. In early ultrastructural studies, no clear junction is identified in these cell types [1–3, 22]. E-Cadherin has been also found to localize outside of the junction, known as non-junctional Cadherin [23–27], suggesting that E-Cadherin may also play a role without forming junctions.

In this study, we show that the loss of E-Cadherin causes unexpected cell-cell fusion between developing germline cysts and surrounding FCs. Our transmission electron microscopy (TEM) study revealed that absence of E-Cadherin leads to the abnormal development of membrane protrusions from both germ cells and FCs in early egg chambers, and a complete absence of plasma membranes at the germline-soma boundary. Consistent with TEM observations, we found mislocalization of germline-specific Vasa protein in FCs, likely caused by diffusion of cytoplasmic contents between germ cells and FCs. FCs receiving germline contents show compromised patterningof FC subpopulations, and the germline cysts or egg chambers show defect in their individualization. Taken together, our results demonstrate that the heterotypic interaction of germline and soma through E-Cadherin is indispensable in maintaining cell-cell boundary integrity between two adjacent tissues.

## Results

### Ultrastructural characterization of germline-soma interaction in the *Drosophila* ovary

We conducted transmission electron microscopy (TEM) analysis for various stages of oogenesis. The germarium is divided into 4 regions (Regions 1, 2a, 2b and 3, Figure 1A) In the anterior compartment (Region1, Figure 1A) of the germarium, containing the niche and early germline cysts and somatic cells, GSCs are known to form adherens junctions with cap cells (CCs), their niche cells [6]. Consistently, we observed junctions present between the GSC and the CC (Figure 1B), suggesting that GSCs form adherens junctions with their niche. These junctions resembled linear adherens junctions (lAJ) or zonula adherens [28], appearing as electron dense areas on the intracellular periphery of the adjacent plasma membranes and a less electron-dense area in the narrow extracellular space between the two cells.

**Figure 1.**
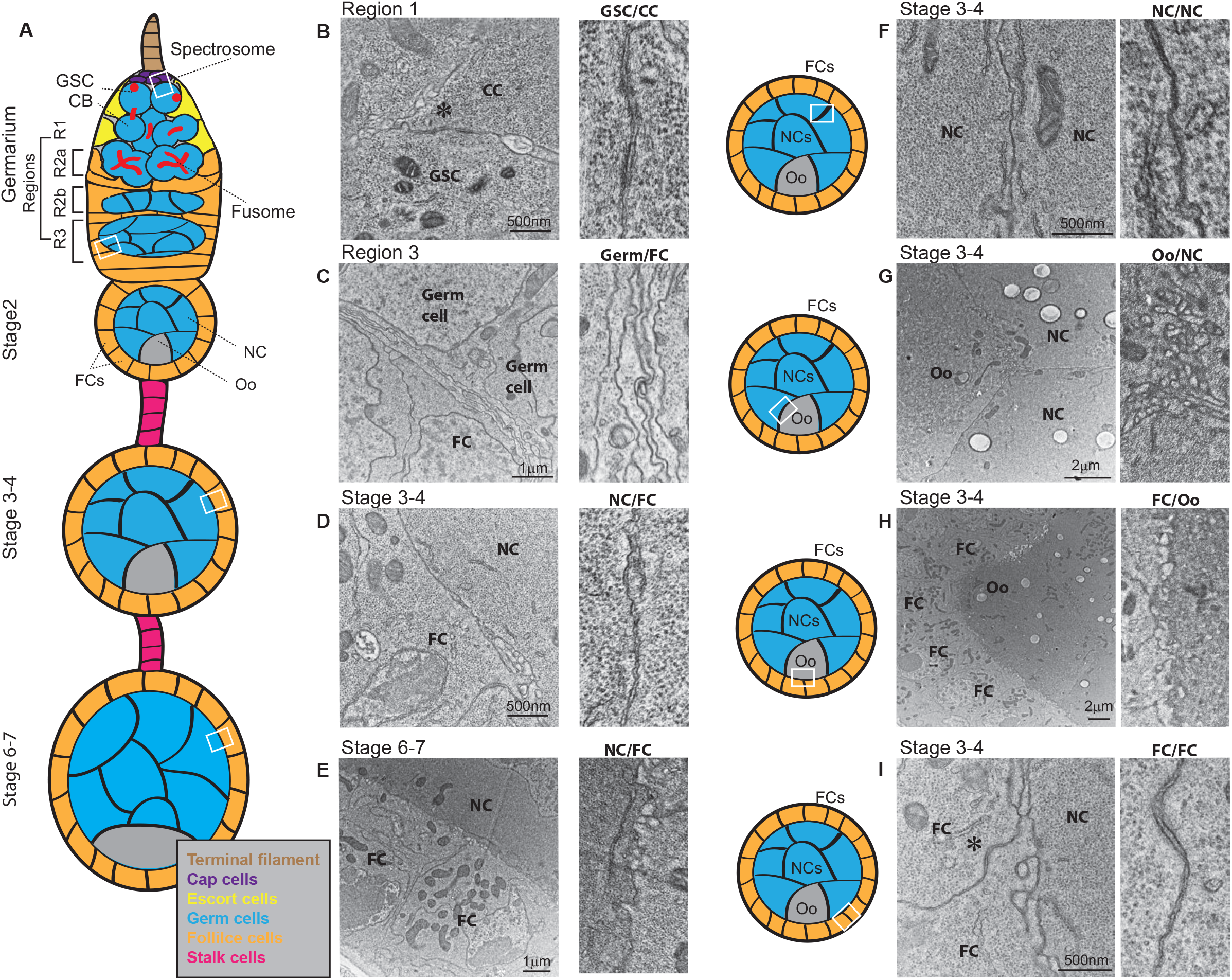
Ultrastructural characterization of germline-soma interaction in the *Drosophila* ovary. **A**) A schematic of an ovariole showing stages of egg chamber development. The germarium houses the germline stem cells (GSCs) which are maintained by associating with their niche (terminal filament; brown and cap cells; purple). Other somatic cells (escort cells; yellow, and follicle cells; orange) support GSC differentiation. GSCs divide asymmetrically to produce one GSC and one cystoblast (CB), which then further divides into syncytial cysts which are encapsulated by follicle cells, forming an egg chamber. As the early cysts divide, the germline-specific organelle, the fusome (red), branches between the interconnect germ cells. Egg chambers released from the germarium are connected by a string of somatic stalk cells; pink. Germ cells are shown in blue. A legend defining the name and color of each cell type is provided at the bottom of the schematic. NC; nurse cell, FCs; follicle cells, Oo; oocyte. **B-I**) Representative transmission electron micrographs of the approximate regions indicated by white boxes in **A**. Magnifications of the cell-cell boundary in the corresponding images are shown in right panels. Asterisks in **B** and **I** indicate adherens junctions. Wildtype (*y,w*) flies were used in 0-4 days post eclosion.

In the posterior compartments (Region 2a/b and 3, Figure 1A) of the germarium, membrane processes from somatic support cells (escort cells: ECs or inner germarial sheath cells: IGS) were observed as a multilayered meshwork between germline cysts. These EC projections were seen tightly wrapping the entire surface of germline cysts (Figure 1C), as previously described [29–33].

We next examined early egg chambers after release from the germarium. These egg chambers contain a germline cyst surrounded by a single layer of FCs, and tight membrane attachments were present between the germline and FCs. Tight alignment of germline and FC membrane was observed throughout the entire germline-FC boundary, closely joining the two tissues with little or no extracellular space (Figure 1D). Tight attachment of germline-FC remained in later egg chambers until approximately stage 6-7, when the FCs and germline cyst project microvilli into a growing extracellular space [4, 34, 35] (Figure 1E). Tight membrane attachment was also observed in NC-NC boundaries and FC-FC boundaries throughout the stages (Figure 1F, G). At all stages analyzed, we observed adherens junctions between FCs (Figure 1G, asterisk) as reported previously [36, 37], and unique membrane processes between oocytes and NCs or between oocytes and FCs (Figure 1H, I) as reported recently [22].

Taken together, these results illustrate the nature of cellular organization by tight adhesion of distinct cell populations, suggesting that the tissue attachment may be occurring in a highly coordinated manner throughout the oogenesis.

### Loss of E-Cadherin results in the formation of abnormal cellular protrusions and cell-cell fusion between germline-FC

In order to characterize function of E-Cadherin in attachment of germline and FCs, we depleted E-Cadherin either in these cell types, by using short-hairpin RNA (shRNA)-mediated knock-down of E-Cadherin. We reasoned that if the germline-FC alignment requires homophilic interaction of E-Cadherin, then depletion of E-cadherin in either tissue should disrupt their organization. To this end, we used the germline-specific *nanos* (*nos*) *Gal4* or the somatic cell driver traffic jam (*tj*) *Gal4* to drive expression of shRNA against *Drosophila E-Cadherin, shg* (*shg^RNAi^*), and analyzed the ultrastructure of early egg chambers. Using a temperature-sensitive allele of *tubGal80* combined with *nosGal4* and *tjGal4* (*nos^ts^* and *tj^ts^*, respectively), we knocked-down *shg* in a tissue-specific and temporally-controlled manner to prevent lethality and to minimize secondary effects (Figure 2A).

**Figure 2.**
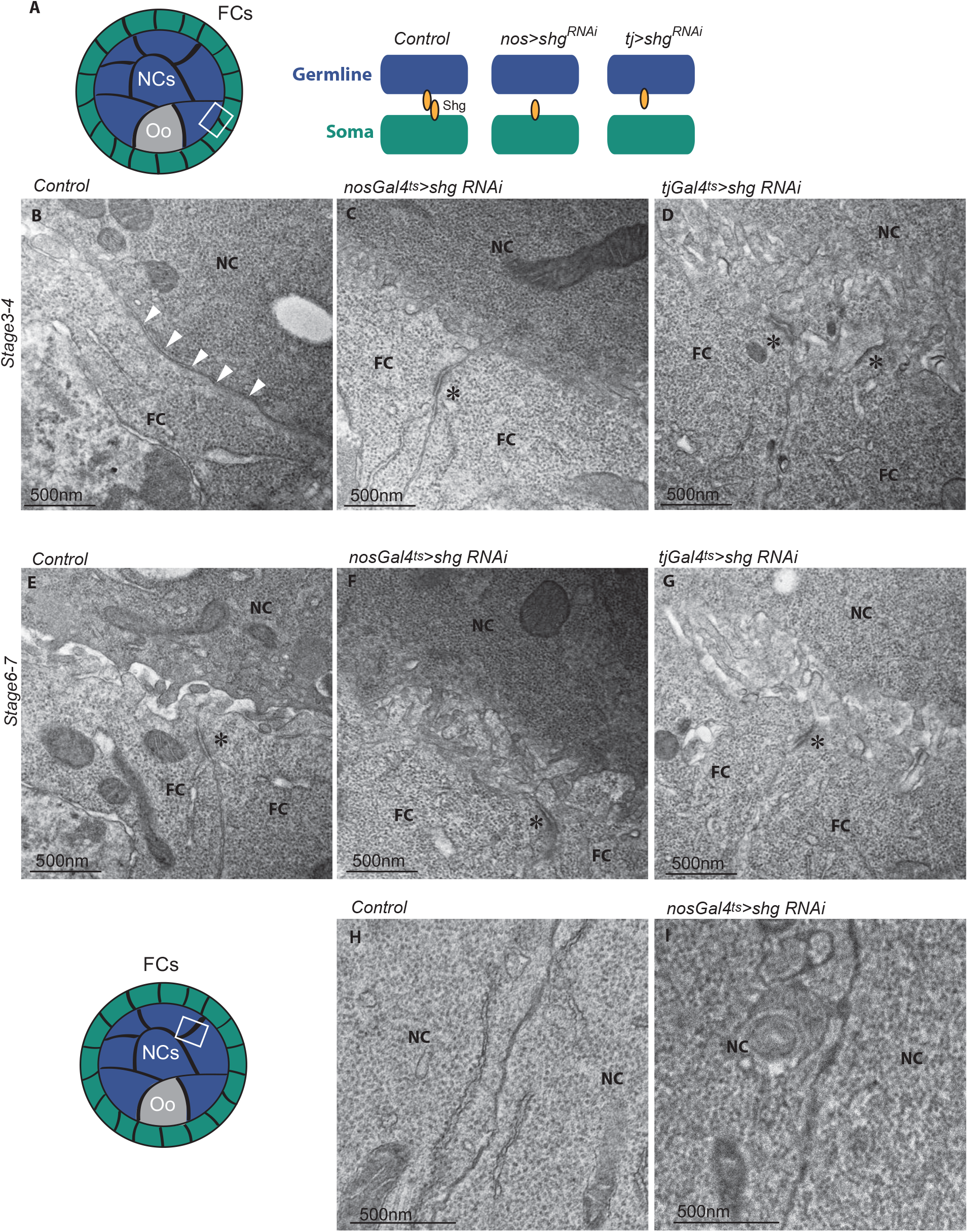
Loss of E-Cadherin results in the formation of abnormal cellular protrusions and cell-cell fusion between germline-FC. **A**) A schematic showing the experimental design of germline and follicle cell knock-down of the gene encoding *Drosophila* E-cadherin, *shotgun (shg*). Flies carrying alleles for temperature-sensitive drivers for either the germline (*nos^ts^*) or follicle cells (*tj^ts^*) were crossed with flies carrying alleles for *shg* shRNA (*shg^RNAi^*; see Methods for a description of genotypes used). Progeny aged 0-7 days post-eclosion were temperature-shifted for 2 days and then dissected and processed for TEM. **B-G**) Representative electron micrographs of regions between the germline and follicle cells (white box shown in **A**) of stage 3-4 (**B-D**) or stage 6-7 (**E-G**) of the indicated genotypes. **H, I**) Representative electron micrographs of regions between the NCs (white box in left diagram) of stage 6-7 of the indicated genotypes. Similar results were obtained by 2 more independent biological replicates. Asterisks indicate adherens junctions.

In wild type and control ovaries, we could clearly trace the plasma membrane boundary at the NC-FC interface, regardless of stages and locations. In early stage of egg-chambers (<Stage6), NC-FC interface were often flat (Figure 2B). Intriguingly, ovaries isolated from both *nos^ts^>shg^RNAi^* and *tj^ts^>shg^RNAi^* showed a complete absence of the flatness of their membrane. Beginning in early egg chambers (stage 3-4), we tended to observe an abundance of cellular protrusions extending from both cell types (Figure 2C, D). In contrast to the membrane protrusions that form in wild type ovaries at stage 6-7 (Figure 2E), these protrusions in *shg^RNAi^* ovaries appeared to integrate into the opposing cell, instead of interdigitating within the extracellular space. A similar pattern was observed throughout various stages (Figure 2F, G). Moreover, the areas where the germline and FCs directly contact one another were completely devoid of plasma membrane, with the cytoplasm of the two cells seemingly continuous (Figure 2D, E). 50% (5 out of 10 ovarioles) of observed early egg-chambers (stage 3-6) were almost completely lacking plasma membrane between FCs and NCs. These results suggest a potential role of E-Cadherin in maintaining the boundary between germline and soma.

It should be noted that the abnormal loss of plasma membrane was also seen in FC-FC boundary in *tj^ts^>shg^RNAi^* (Figure 2D, G), while there were no detectable changes in germline-germline (NC-NC) boundaries in *nos^ts^>shg^RNAi^* (Figure 2H-I).

### Loss of E-Cadherin results in cytoplasmic leakage between FC and germline cysts and progressive individualization defect

We next addressed the consequences of a disrupted barrier between germline and soma during egg chamber development. We reasoned that if the FCs and germline fused, then cytoplasmic contents between the two should be shared and therefore could be visualized by immunofluorescence staining. Indeed, staining for the germline-specific protein Vasa revealed that some FCs of *shg^RNAi^* ovaries contained Vasa (Figure 3B-G). Similar phenotype was observed by using a different *shg^RNAi^* construct, confirming that the phenotype was specifically caused by loss of *shg* gene product (Figure S1A). Importantly, the Vasa-positive FCs of *shg^RNAi^* were not observed as frequently as cellcell fusion observed in TEM. We observed FC-germline fusion in approximately 50% of ovarioles in 7 day-old females (Figure 3J), suggesting that the boundary of FC/germline may still maintain some diffusion barrier. Indeed, the difference of electron density between NC and FC was still maintained especially in *nosGal4* driven *shg^RNAi^* samples (Figure 2C, F). Interestingly. the frequency of ovarioles containing Vasa-positive FCs significantly increased with the age of the animal when expressed in the germline (by *nos^ts^*) or FCs (by *tj^ts^*) (Figure 3J), suggesting that unknown age-related factors may play a role in preventing the diffusion of germline contents.

**Figure 3.**
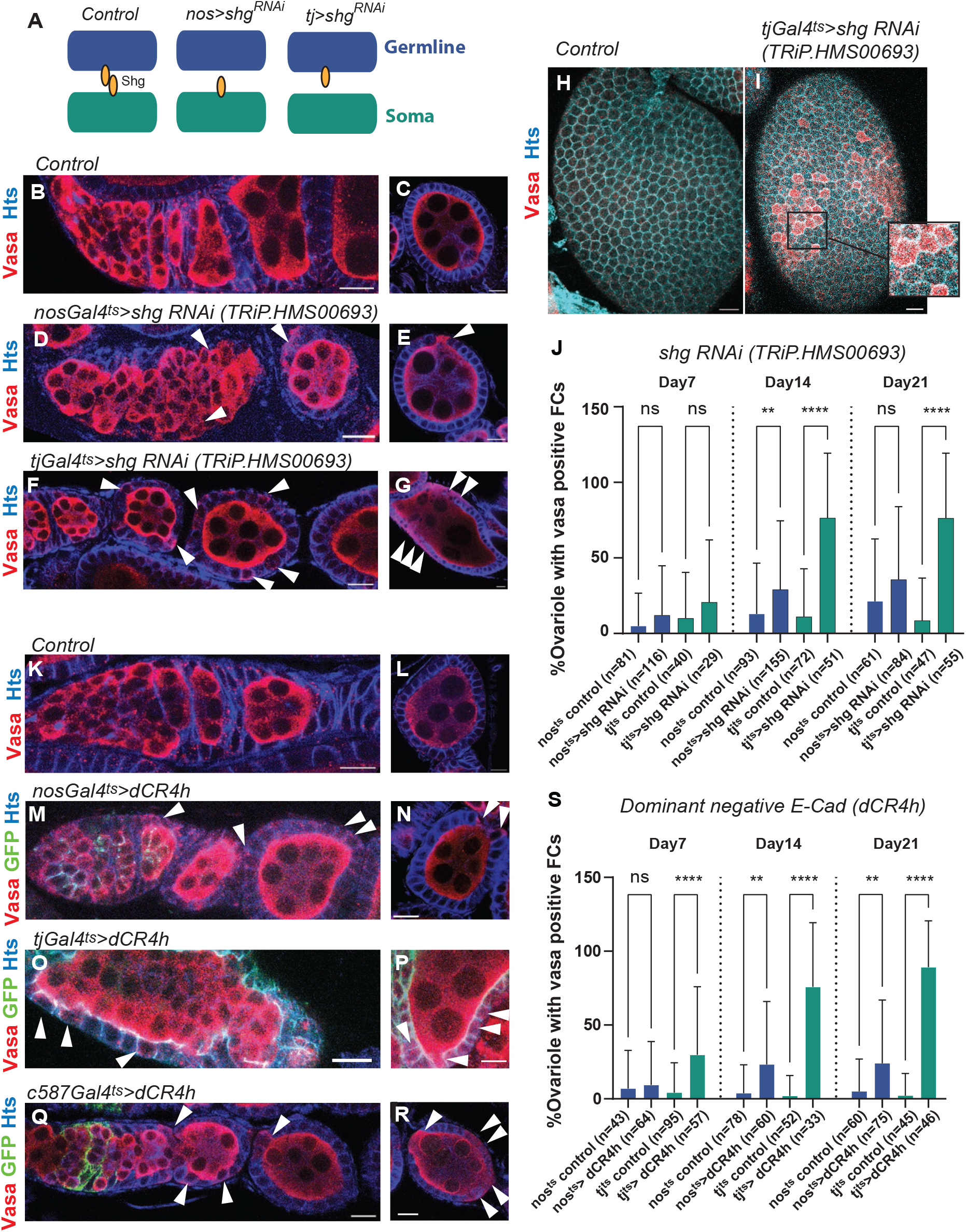
Loss of E-Cadherin-based attachment results in cytoplasmic leakage between FC and germline cysts. **A**) A schematic showing the experimental design of germline and follicle cell knock-down of the gene encoding *Drosophila* E-cadherin, *shotgun* (*shg*). **B-G**) Representative immunofluorescent (IF) staining images. Germline (*nosGal4^ts^*) (**D, E**) or follicle cell (*tjGal4^ts^*) (**F, G**) were used to knockdown *shg* using indicated RNAi lines. Samples were used after 7-days temperature shift for all images. Left panels (**B, D, F**) show germaria to stage 3-4 egg chambers. Right panels (**C, E, G**) show stage 6-7 egg chambers. White arrowheads indicate abnormal localization of Vasa in the FC layer. **H, I**) Surface views of stage 6-7 egg chambers of indicated genotypes. The inset in **I** shows Vasa positive FCs. **J**) A graph shows frequency of ovarioles containing Vasa-positive FCs in indicated genotypes after indicated days of temperature shift. **K-R**) Representative IF staining images. Germline (*nosGal4^ts^*) (**M, N**), follicle cells (*tjGal4^ts^*) (**O, P**) or escort cells (*c587Gal4^ts^*) drivers were used to express dominant negative form of E-Cadherin (*dCR4h*). Samples were used after 7-days temperature shift for all images. Left panels (**K, M, O, R**) show germaria to stage 3-4 egg chambers. Right panels (**L, N, P, R**) show stage 6-7 egg chambers. White arrowheads indicate abnormal localization of Vasa in the FC layer. Green signal in **M-R** indicates dCR4h expression. **S**) A graph shows frequency of ovarioles containing Vasa-positive FCs in indicated genotypes after indicated days of temperature shift. For **J** and **S**, driver only flies were used for control. Adjusted p-values from Šidák’s multiple comparisons test are shown as *p<0.05, **p<0.01, ***p<0.001, ****p<0.0001; ns, non-significant (p≥0.05). n: number of scored ovarioles. Similar results were obtained by 2 more independent biological replicates. Scale Bars: 10 μm.

So far, the emergence of Vasa-positive FCs together with fusion between FC and germline cysts observed in TEM strongly suggests that loss of E-Cadherin leads to disruption of the physical barrier between the germline and FCs resulting in “leakage” of germline cytoplasmic contents into neighboring FCs. However, there is the possibility that FCs lacking E-Cadherin may alter intracellular signaling in FCs, which leads to expression of germline genes in FCs. To distinguish between these possibilities, we took advantage of a dominant-negative form of E-cadherin-GFP (dCR4h) that retains the transmembrane and intracellular domains but lacks part of the extracellular domain so that homophilic interactions are abolished without affecting intracellular signaling [38].

Overexpression of a dominant negative E-cadherin, dCR4h, with either *nos^ts^* or *tj^ts^* resulted in the emergence of Vasa-positive FCs (Figure 3K-P), confirming that Vasa-positive FCs result from a loss of physical adhesion of membranes mediated by homophilic interaction of E-Cadherin between the germline and FCs and not by altering intracellular signaling by the intracellular domain of E-Cadherin.

Importantly, Vasa leak was observed throughout later stage egg chambers even when dCR4h-GFP expression was only expressed in the germarium, by early germline under the *nosGal4* driver (Figure 3M, N, Figure S1B), or in early somatic cells under the *c587Gal4* driver (Figure 3Q, R), suggesting that the E-Cadherin-based interaction within the germarium is indispensable for priming germline-soma interaction in later stages.

Previous reports have demonstrated that the misexpression or mislocalization of germline gene products in FCs results in cyst or egg-chamber individualization defect, where two or more germline cysts are surrounded by a single FC epithelium [39] [40]. FCs expressing germline determinants were thought to alter their cell fate and fail to properly specify FC subtypes [40]. Consistent with these reports, we found that *shg^RNAi^* and *dCR4h* ovaries exhibit fused egg chambers accompanied with ectopic expression of laminC, the marker of stalk cells, in main body FCs (Figure 4, Figure S2).

**Figure 4.**
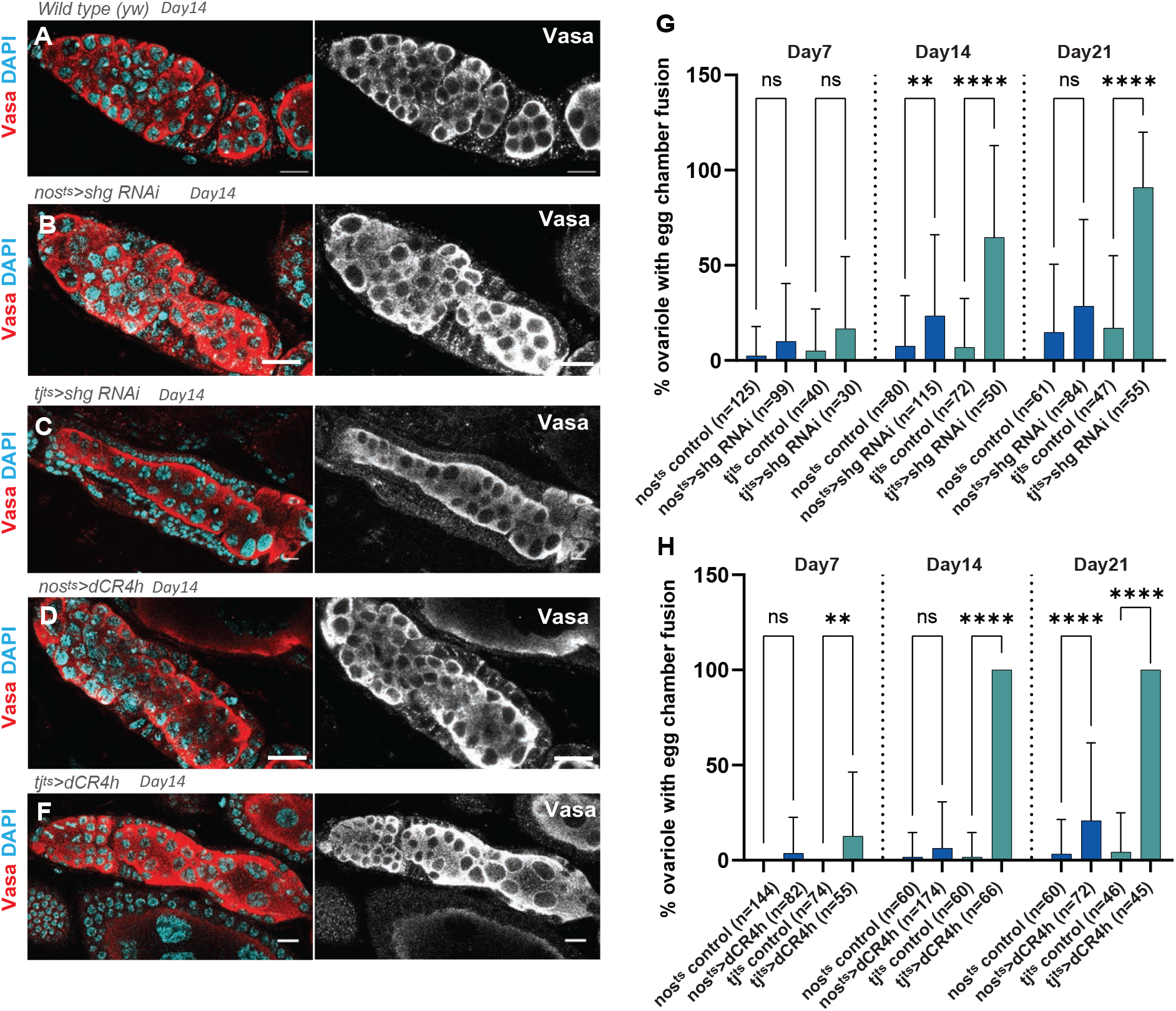
Loss of E-Cadherin-based attachment results in progressive fusion of egg chambers. **A-F**) Representative IF staining images. Germline (*nos^ts^*) or follicle cell (*tj^ts^*) were used to knockdown *shg (TRiP.HMS00693*) or expression of dCR4h. Samples were used after 14-days temperature shift for all images. Right panels show Vasa channel only. **G, H**) Graphs show frequency of ovarioles containing fused egg chambers in indicated genotypes after indicated days of temperature shift. Adjusted p-values from Šidák’s multiple comparisons test are shown as *p<0.05, **p<0.01, ***p<0.001, ****p<0.0001; ns, non-significant (p≥0.05). Similar results were obtained by 2 more independent biological replicates. n: number of scored ovarioles. Scale Bars: 10 μm.

Taken together, our results suggest a role for E-Cadherin-based interaction of germline cysts and FC epithelia that is required for maintaining the distinct boundary between the two cell types. A disruption of this boundary results in progressive cytoplasmic leak between adjacent cells and egg chamber fusion.

## Discussion

E-Cadherin plays a role in an adhesion between cells and is essential for various tissue functions and development. In this work, we show that homophilic interaction of E-Cadherin expressed in germline and soma is required for proper interaction of these cell types. From the germarium to early egg chamber stages, developing germ cells and surrounding somatic cells tightly adhere to each other. We found that loss of E-Cadherin not only results in loosened attachment, but also results in extensive fusion of these cell types. The aberrant cell-cell fusion, in turn, causes mislocalization of germline components into somatic cells and subsequent egg chamber fusion, suggesting a crucial physiological function of E-Cadherin-based cell-cell interaction between two adjacent tissues.

It is interesting to note that the lack of germline-soma junctions results not only in tissue architecture defects, but in an apparent dissolution of the plasma membranes of interacting two cell types, such that the boundary between the two is indistinguishable.

This observation is reminiscent of physiological cellular fusion seen in other systems. For example, across several species, the fusion of myoblasts involves the fusion of adjacent cell membranes and is essential for the development of mature muscle tissue [41,42]. Additionally, the fusion of sperm and egg likewise involves the fusion of plasma membranes of two distinct cells, and its mechanisms have puzzled researchers for decades [43–45]. Intriguingly, in both cases, membrane protrusions are associated with fusion as a possible prerequisite. For myoblast fusion, the apposition of two adjacent membranes is facilitated by the formation of F-actin protrusions, followed by the formation of fusion pores which ultimately fuse the two cells [41,42, 46]. Formation of protrusive membrane structures is also observed in sperm-oocyte fusion and is essential for successful fertilization [44, 45, 47].

Involvement of E-Cadherin in cell-cell fusion has been suggested in a few systems, including sperm-oocyte fusion [48], macrophage and T-Cell fusion [49], and *Drosophila* muscle cell fusion [50], suggesting the possibility that the *Drosophila* egg chamber could be a new model to study impaired cellular fusion and to address its effect on many physiological and pathological conditions. Investigating the potential interplay between homophilic interaction of E-Cadherin, cellular protrusions, and cellcell fusion would likely be a fascinating future study.

## Figure Legends

**Figure S1.**
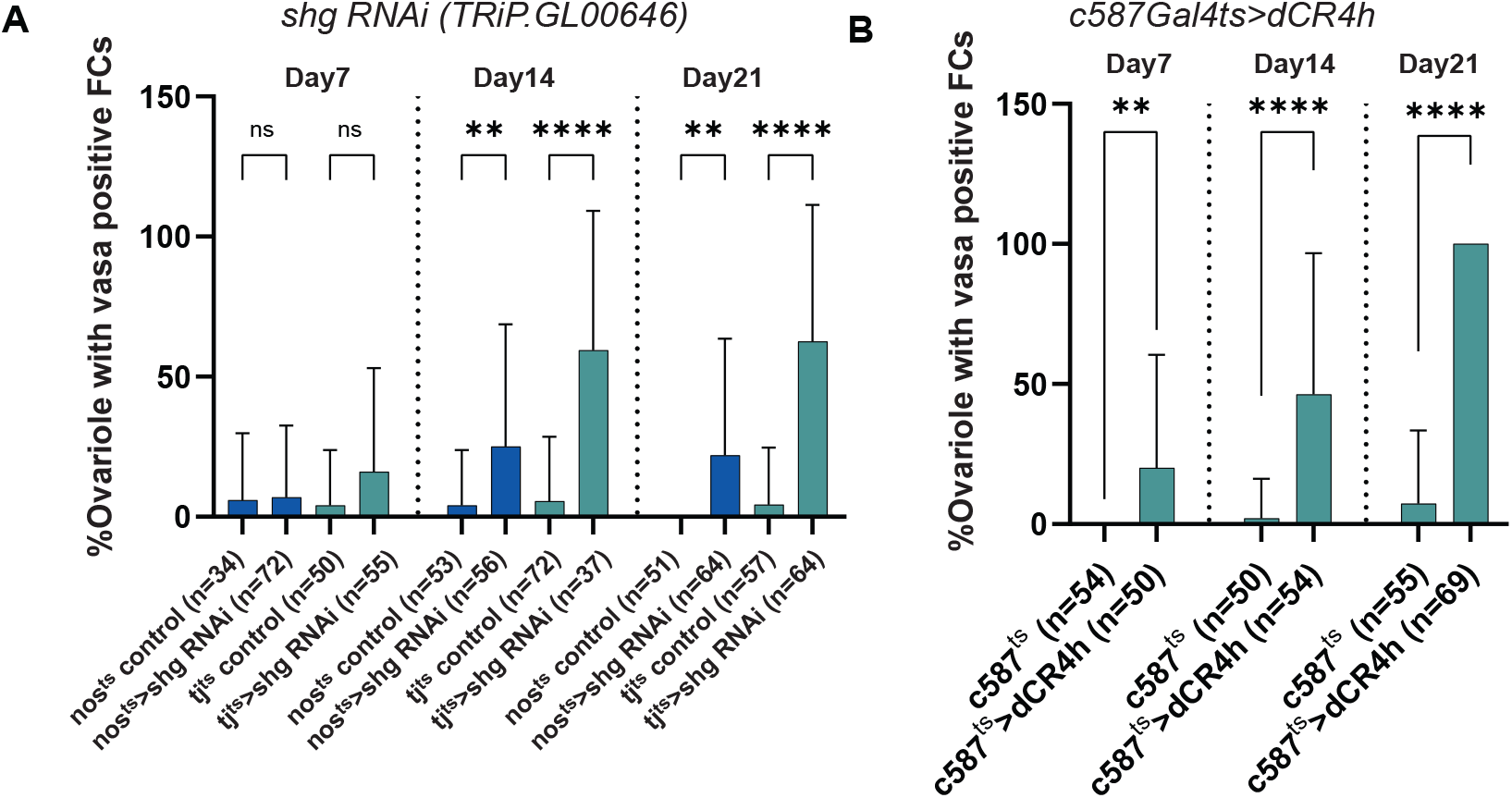
**A-B**) Graphs show frequency of ovarioles containing Vasa-positive FCs in indicated genotypes after indicated days of temperature shift. Adjusted p-values from Šidák’s multiple comparisons test are shown as *p<0.05, **p<0.01, ***p<0.001, ****p<0.0001; ns, non-significant (p≥0.05). Similar results were obtained by 2 more independent biological replicates. n: number of scored ovarioles.

**Figure S2.** **A-F**) Representative IF staining images. Germline (*nos^ts^*) or follicle cell (*tj^ts^*) were used to knockdown *shg (TRiP.HMS00693*) or expression of dCR4h. Samples were used after 7-days temperature shift for all images. **G**) Quantification of the frequency of ovarioles showing stalk cell mislocalization. Adjusted p-values from Šidák’s multiple comparisons test are shown as *p<0.05, **p<0.01, ***p<0.001, ****p<0.0001; ns, non-significant (p≥0.05). Similar results were obtained by 2 more independent biological replicates. n: number of scored ovarioles. Scale Bars: 10 μm.

## Materials and methods

### Fly husbandry and strains

All fly stocks were raised on standard Bloomington medium at 25°C (unless temperature control was required), and indicated age (days after eclosion) of flies were used for all experiments. For temperature shift experiments, *nosGal4* (delta VP16), *tjGal4* or *c587Gal4* flies were used combined with *tubGal80^ts^* to induce short hairpin RNA expression or dCR4h for indicated days at 29°C. Stocks used were *nosGal4 delta VP16* [51]; *tubGal80^ts^* [52], *tj-Gal4* (FBtp0089190), and *UAS-dCR4h* (FBal0103562) [53] (gifts from Yukiko Yamashita). Stocks obtained from the Bloomington Drosophila Stock Center were: *y,w* (Stock #189, BDSC) and *shg^RNAi^* TRiP.HMS00693 and TRiP.GL00646 (BDSC).

### Transmission electron microscopy

Ovaries of the indicated genotype were dissected and processed as described previously [54]. Briefly, ovaries were dissected into phosphate buffered saline (PBS) and then fixed in a solution of 2.5% glutaraldehyde and 3% paraformaldehyde in 0.1 M sodium cacodylate buffer on ice. Samples were then washed in cacodylate buffer containing 2 mM calcium chloride and incubated in a solution of 1.5% potassium ferrocyanide and 2% osmium tetroxide in in cacodylate buffer, followed by washing with water and a subsequent incubation in 2% aqueous osmium tetroxide at room temperature. Samples were then washed with water and placed in 1% aqueous uranyl acetate overnight at 4°C.

The next day, samples were then dehydrated via a graded series of alcohol dilutions, then washed with propylene oxide and in epoxy resin and allowed to polymerize at 60°C for 48 hours.

Ultrathin sections (60 nm) of Lowicryl HM-20-embedded follicles were cut on a UC-7 ultramicrotome (Leica Biosystems) with a diamond knife (Diatome, Hatfield, PA) and imaged using Hitachi H-7650 transmission electron microscope.

### Immunofluorescence staining

Ovaries were dissected into 1X phosphate-buffered saline (PBS) and fixed in 1 ml of 4% formaldehyde in PBS for 30-60 minutes, then washed three times in 1ml of PBS + 0.3% TritonX-100 (PBST) for one hour, then incubated in primary antibodies in 100μl of 3% bovine serum albumin (BSA) in PBST at 4°C overnight. Samples were then washed three times in 1 ml of PBST for one hour (three 20-minute washes), then incubated in secondary antibodies in 100 μl of 3% BSA in PBST for 2-4 hours at room temperature, or at 4°C overnight. Samples were then washed three times in 1 ml of PBST for one hour (three 20-minute washes), then mounted in a drop of VECTASHIELD with 4,6-diamidino-2-phenylindole (DAPI) (Vector Lab, H-1200).

Primary antibodies used were: rat anti-Vasa (1:20), mouse anti-Hts (1B1, 1:20), mouse anti-LaminC (1:20) from Developmental Studies Hybridoma Bank (DSHB). Secondary antibodies used were Alexa Fluor 568-Goat Anti-Rabbit IgG H&L (1:400, Abcam, ab175471), Cy5-Goat Anti-Rat IgG, Cy3-H&L Goat Anti-Mouse IgG H&L (1:400, Jackson Immuno Research Labs).

### Phenotype analysis

The “vasa positive follicle cell” phenotype was determined by the presence of Vasa in one or more follicle cells in an egg chamber, as seen by immunofluorescence staining with Vasa, and DAPI. The “egg chamber fusion” phenotype was determined as egg chambers containing more than 16 germ cells encapsulated by a single layer of follicle cells, as seen by immunofluorescence staining with Vasa and DAPI.

To quantify the frequency of phenotypes, ovarioles were scored using a YES/NO criteria. Ovarioles from a minimum of ten ovaries were dissected from flies in the same culture and scored as containing one or more egg chambers exhibiting the phenotype of interest (“YES”), or no egg chambers exhibiting the phenotype of interest (“NO”), and the total number of ovarioles with the phenotype was calculated as a frequency (percentage). All data are shown as means ± s.d.

### Statistical analysis and graphing

No statistical methods were used to predetermine sample size. The experiment values were not randomized. The investigators were not blinded to allocation during experiments and outcome assessment. Statistical analysis and graphing were performed using GraphPad prism9. All data are shown as means ± s.d. The adjusted P values from Šidák’s multiple comparisons test are provided; shown as *P<0.05, **P<0.01, ***P<0.001, ****P<0.0001; NS, non-significant (P≥0.05).

## Acknowledgements

We thank Yukiko M. Yamashita and the Bloomington *Drosophila* Stock Center and the Developmental Studies Hybridoma Bank for reagents. This research is supported by R35GM128678 from National Institute for General Medical Sciences and start-up funds from UConn Health (to M.I.).

## Author Contributions

M.I, and M.A; conception and design, acquisition of data, analysis and interpretation of data, R.N; Implementation of TEM analysis, interpretation of data. All authors wrote and edited the manuscript.

